# *Rheinheimera palamurensis* sp. a novel bacterium isolated from drinking water resources

**DOI:** 10.1101/2025.04.03.647078

**Authors:** T. Ranjith Paul, A Shiva Shanker, K. Sanjeev Kumar, Dasaiah Srinivasulu, Praveen Kumar Vootla, Pavan Kumar Pindi

**Affiliations:** Department of Microbiology, Palamuru University, Mahabubnagar –509001, Telangana State, India

**Keywords:** *Rheinheimera palamurensis* sp. nov., *Gammaproteobacteria*, *Chromatiales*, *Chromatiaceae*, Polyphasic taxonomy

## Abstract

A bacterial strain designated NG3^T^ was isolated from a drinking water samples collected from Nagar Kurnool, Telangana State, India (16° 29’ 14.5356’’ N ; 78° 18’ 33.966’’) and characterized using a polyphasic taxonomic approach, Strain NG3^T^ was Gram-negative, aerobic, non pigmented and motile by means of a single polar flagellum. Growth occurred at 18 - 37° C (optimum 27 - 32° C), pH at 6.5 – 8.0 (Optimum 7.0 – 7.5) and in the presence of 0-2% NaCl (Optimum 0.7-1.0%), Phylogenetic analyses based on 16S rRNA gene sequence showed that strain NG3^T^ belonged to the genus Rheincheimera and its most closely related neighbor was *Rheinheimera aquatica* GR5^T^ (GQ168584) with sequence similarity of 99 %. The cellular fatty acid composition of strain NG3^T^ showed a spectrum of 11 fatty acids with a pronounced dominance of saturated fatty acids (82.7 %), including a high abundance of C16 : 0, C18 : 1ω7c, and C12 : 0 3-OH. The major respiratory quinone was MK-7 and Q-8. The DNA G+C content of the genomic DNA was 48.5 mol%. The polar lipid profile consisted of a mixture of phosphatidylethanolamine, phosphatidylglycerol, phosphatidylserine, aminolipid and two uncharacterized phospholipids. The DNA–DNA relatedness of strain NG3^T^ with respect to recognized members of the genus *Rheinheimera* was less than 70 %. On the basis of the genotypic, chemotaxonomic and phenotypic data, strain NG3^T^ represents a novel species in the genus *Rheinheimera*, for which the name *Rheinheimera palamurensis* sp. nov. was proposed. The type strain is NG3^T^ (=GU566360=JCM16716=KCTC23111).

## Introduction

The genus *Rheinheimera*, a member of the family *Chromatiaceae* of the class *Gamma-proteobacteria*, was first proposed with *Rheinheimera baltica* as the type species by Brettar *et al*. 2002 and later amended by Merchant et al. (2007), Chen et al. (2010), Li et al. (2011), Liu et al. (2012), Hayashi et al. (2018) and Sheu et al. (2018). The members of the genus *Rheinheimera* shared some common features, being Gram-stain-negative, aerobic, motile and oxidase- and catalase-positive. At the time of writing this manuscript, the *Rheinheimera* genus comprised of 24 species with validly published names. Members of *Rheinheimera* genus are known for their diverse habitats such as fresh water, fishing hooks, coastal sediment, seawater, seashore sand, sediments, the central Baltic Sea, marine sediment, chironomid egg mass, soil, rice roots and deep-sea water as per Hayashi et al. (2018). Herein, we report the taxonomic characteristics of a novel species of the genus *Rheinheimera* isolated from drinking water sample collected from Nagar Kurnool, Telangana State, India.

## Materials and Methods

### Isolation and Bacterial culture conditions

Strain NG3^T^ was isolated from a drinking water sample collected from Nagar Kurnool, Telangana State, India (16° 29’ 14.5356’’ N ; 78° 18’ 33.966’’) on 20^th^ March, 2010. The drinking water sample that yielded strain NG3^T^ had a pH of 7. For isolation of bacteria, 100 μl of water sample was placed on nutrient agar medium adjusted to pH 6.5–8 with salt percent ranging from 0–2 %, and incubated at room temperature for 7 days. Based on the colony morphology, a whitish colony was selected and characterized in the present study. Purification of the isolate was achieved by streaking on tryptone soy agar (TSA) plates repeatedly.

### Morphological, Physiological and biochemical characteristics

Cell morphology and motility were studied using a light microscope. Motility was assessed on TSA medium containing (l–1): pancreatic digest of casein (17 g), papaic digest of soyabean meat (3 g), sodium chloride (5 g), dipotassium hydrogen phosphate (2.5 g), dextrose (2.5 g) and agar (0.4 g). Growth on TSA medium at 18-37°C temperatures, salt tolerance, biochemical characteristics, carbon assimilation, H_2_S production and the sensitivity of the culture to different antibiotics were determined by previously described methods (Lanyi, 1987; Smibert & Krieg, 1994). Biochemical characteristics were also double-checked with Hi25 Enterobacteriaceae identification kit (cat. #KB003; Hi Media) and Hi Carbohydrate kit parts A, B and C (cat. # KB009; Hi Media) according to the manufacturer’s protocol. Growth of strain NG3^T^ at different pH was checked on NA broth buffered either with citric acid-NaOH (for pH 5 and 6) and phosphate (for pH 7 and 8).

Fatty acid methyl esters were prepared and analysed by using the Sherlock Microbial Identification System (MIDI) according to the protocol described by Agilent Technologies. For this purpose, all strains were grown on TSA medium at 30 °C for 2 days. Peptidoglycan was prepared according to the method described by Rosenthal & Dziarski (1994). The peptidoglycan obtained was then hydrolysed with 2 M HCl at 60 min. The hydrolysate was extracted into acetate buffer and subjected to amino acid analysis on an automated amino acid analyser. The composition of the peptidoglycan was determined according to the method of Schleifer & Kandler (1972). Polar lipids were analysed by lyophilizing cell pellets free of medium by the method of Minnikin et al. (1975) and identified by TLC. Cell-wall components were extracted and analyzed according to the method described by Komagata & Suzuki (1987). Menaquinones and polar lipids were determined in freeze-dried cells. Menaquinones were extracted as described by Collins et al. (1977) and were analysed by HPLC (Groth et al., 1997).

### Phylogenetic Analysis

DNA was isolated according to the procedure of Marmur (1961) and the G+C content was determined from melting point (Tm) curves (Sly et al., 1986) obtained by using a Lambda2 UV-Vis spectrophotometer (Perkin Elmer) equipped with the Templab 2.0 software package (Perkin Elmer). *Escherichia coli* K-12 strain was used as a standard in determining the mol% G+C content of the DNA. For 16S rRNA gene sequencing, DNA was prepared using the Mo Bio microbial DNA isolation kit (Mo Bio Laboratories) and sequenced as described previously (Lane, 1991). Taq polymerase supplied by Merck was used for PCR, which was started with the primers 16SF 5′-GTTTGATCCTGGCTCAG-3′ and 16SR 5′ AAGGAGGTGATCCAGCCGCA-3′. The resultant almost-complete sequence of the 16S rRNA gene of strain NG3^T^ contained 1495 nucleotides. This sequence was subjected to BLAST sequence similarity search (Altschul et al., 1990) and analysis using the EzTaxon server (Kim et al., 2012) to identify the nearest taxa. All the 16S rRNA gene sequences belonging to the family Planococcaceae were downloaded from the GenBank database (http://www.ncbi.nlm.nih.gov), aligned using the CLUSTAL X program (Thompson et al., 1997) and the alignment corrected manually. Phylogenetic trees were reconstructed using two tree-making algorithms, maximum-likelihood (ML) using the PhyML program (Guindon & Gascuel, 2003) and neighbour-joining (NJ) (Saitou & Nei, 1987) using the PHYLIP package, version 3.5 (Felsenstein, 1993), and the resultant tree topologies were evaluated by bootstrap analysis based on 1000 resamplings using the SEQBOOT and CONSENSE programs in the PHYLIP package. Pair-wise evolutionary distances were computed using the DNADIST program with the Kimura 2-parameter model as developed by Kimura (1980).

### DNA-DNA Hybridization

The genomic relationship between strain NG3^T^, *Rheinheimera aquatica* GR5^T^ (GQ168584), *Rheinheimera texasensis* A62-14B^T^ (AY701891) and *Rheinheimera chironomi* K19414^T^ (DQ298025) was examined by DNA–DNA hybridization using a membrane filter technique (Tourova & Antonov, 1988). Hybridization was performed with three replications for each sample. A nick translation kit was used for labeling the probe with a-P32 dCTP. The DNA immobilized on the blots (nylon membranes) was probed with labeled DNA and then exposed to a phosphor imaging screen (Amersham Biosciences). A TYPHOON (3480) variable mode imager was used to scan and quantify the phosphor-imaging screen.

### Whole Genome Analysis

#### Read quality check

Checked the following parameters from the fastq file - Base quality score distribution, sequence quality score distribution, average base content per read, GC distribution inthereads, PCR amplification issue, overrepresented sequences and adapters.

Based on the quality report of fastq files, sequences were trimmed to only retain high-qualitysequences for further analysis. In addition, the low-quality sequence reads were excludedfrom the analysis. The adapter trimming was performed using fastq mcf (v-1.04.803).

### Alignment to bacteria, virus, fungi, and archaea genome

The adapter trimmed reads were aligned to the reference genome of bacteria, virus and archaea to check the alignment percentage using BWA (0.7.12).

#### De novo assembly

Denovo assembly was carried out using megahit assembler (v1.2). Megahit makes useof succinct de Bruijn graphs which are compressed representations of de Bruijn graphs. Theassembled scaffolds were taken for further downstream analysis. Blast was performedfor the scaffolds fasta to predict the predominant species.

### Reference Guided assembly

The human unaligned reads were mapped to the reference of the predominant specieswhich was predicted based on the blast results to get the coverage and assembled fasta.

### Variant Prediction and annotation

The Reference aligned reads were used to predict variants using GATK and the variants were annotated using the snpEff. Genome annotation and functional annotation The Primarily assembled genome was annotated using Prokka to predict genes, CDSetc. The functional annotation was performed using Emapper.

### Data Summary

The sample was taken for bacterial whole genome analysis and the data size of the sampleis 4.44 Gb. Q30% was above 93% and the average GC% was above 46%. Kindly refer to the QC report for further details. The quality check for the sample after adapter trimming was performed and the adapter trimmed sequences were processed for down stream analysis.

## Results and Discussion

Cells of strain NG3^T^ were Gram-negative rods of 0.7–0.9 μm in width and 1.1–2 μm in length, and showed binary fission (Fig. 1). Colonies were circular convex, 1–2 mm in diameter, smooth, non-pigmented, opaque form and entire on nutrient agar. The strain was aerobic and showed chemo-heterotrophic growth. Strain NG3^T^ was able to grow at temperature range 18-37 °C (optimum 27-32 °C), pH 6.5-8.0 (optimum 7.0–7.5) with 0 to 2% (w/v) NaCl (Optimum 0.7–1%).

**Figure 1.**
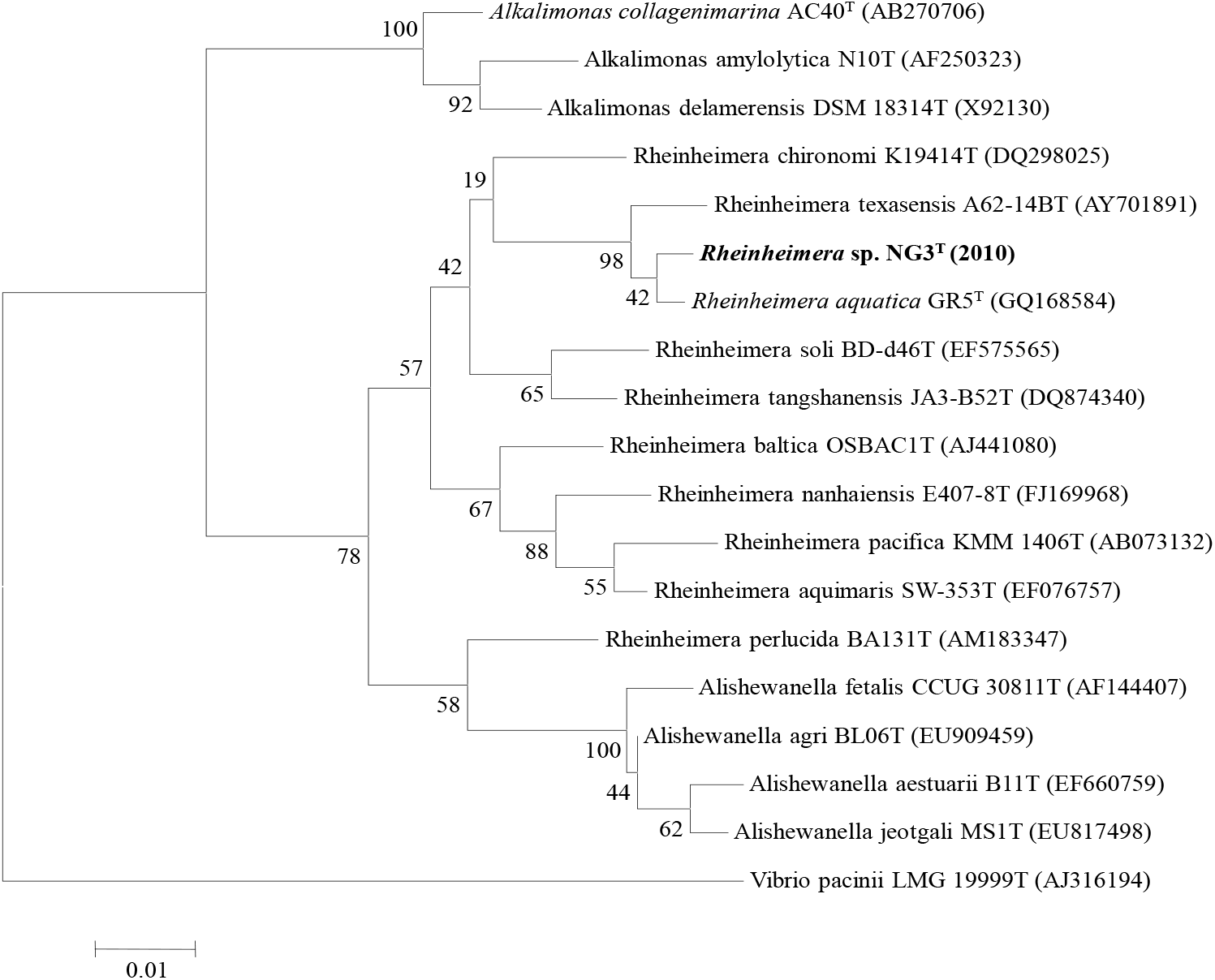
Phylogenetic tree of Rheinhemera sp. NG3(GU566360)

The chemotaxonomic analysis of lipids of the strain NG3^T^ showed the presence of 11 fatty acids with a pronounced dominance of saturated fatty acids (82.7 %), including a high abundance of C16 : 0, C18 : 1ω7c, and C12 : 0 3-OH (Table 1). When compared with *Rheinheimera aquatica* GR5^T^ (GQ168584), *Rheinheimera texasensis* A62-14B^T^(AY7018891), *Rheinheimera chironomi* K19414^T^ (DQ298025), *Rheinheimera soli* BD-d46^T^ (EF575565) and *Rheinheimera tangshanensis* JA3-B52^T^ (DQ874340), the number of fatty acids detected in strain NG3^T^ was lower but the composition differed considerably (Table 1). The predominant quinones present in strain NG3^T^ were MK-7 and Q-8. Strain NG3^T^ exhibited a complex lipid profile consisting of diphosphatidylglycerol, phosphatidylglycerol and phosphatidylethanolamine. The cell-wall peptidoglycan contained mesodiaminopimelic acid as the diamino acid and cell-wall sugars were D-glucose and D-galactose. Other characteristics are listed in the species description. The DNA G+C content of strain NG3^T^ is 48.5 mol%. Based on comparison of several phenotypic features, the strain NG3^T^ could be differentiated from most closely related type species of *Rheinheimera*.

**Table 1.** Cellular fatty acid composition of R. palamurensis NG3T and members of related Rheinheimera species. **Strains: 1, *Rheinheimera aquatica* GR5^T^ (GQ168584); 2, *Rheinheimera texasensis* A62-14BT(AY701891); 3, *Rheinheimera chironomi* K19414T (DQ298025); 4 *Rheinheimera soli* BD-d46T (EF575565); 5, *Rheinheimera tangshanensis* JA3-B52T (DQ874340).**

| Fatty acids | *Rheinheimera*<br>sp. NG3T<br>(GU566360) | *Rheinheimera*<br>*aquatica*<br>GR5<sup>T</sup><br>(GQ168584) | *Rheinheimera*<br>*coerulea*<br>TAPG2 | *Rheinheimera*<br>*texasensis*<br>A62-14BT<br>(AY701891) | *Rheinheimera*<br>*chironomi*<br>K19414T<br>(DQ298025) | *Rheinheimera*<br>*soli*<br>BD-d46T<br>(EF575565) | *Rheinheimera*<br>*tangshanensis*<br>JA3-B52T<br>(DQ874340) |
| --- | --- | --- | --- | --- | --- | --- | --- |
| C10 : 0 | 2.7 | 1.5 | - | 1.2 | 3.2 | 3.0 | 2.9 |
| C11 : 0<br>3-OH | 3.1 | 1.9 | 2.8 | - | 1.0 | - | 2.7 |
| C12 : 0 | 1.3 | 2.8 | 2.8 | 3.1 | 2.9 | 1.0 | - |
| C12 : 0<br>3-OH | 7.2 | 12.0 | 8.8 | 10.0 | 20.1 | 8.5 | 10.0 |
| C14 : 0 | 2.3 | 2.0 | 1.9 | 3.3 | 3.8 | 2.1 | 1.0 |
| C14 : 0<br>3-OH | - | - | 2.8 | - | - | 1.6 | - |
| C15 : 0 | - | - | - | - | - | 1.7 | - |
| C15 :<br>1ω8c | 1.0 | 1.7 | 2.6 | - | 1.0 | - | 3.1 |
| iso-C16 :<br>0 | - | 1.1 | - | - | - | - | 1.1 |
| C16 : 0 | 18.3 | 16.3 | 14.4 | 25.5 | 18.5 | 23.0 | 15.5 |
| C16 :<br>1ω9c | - | - | - | - | - | 1.7 | - |
| C17 : 0 | 1.7 | 1.7 | 2.7 | - | - | - | 2.5 |
| C17 :<br>1ω8c | 5.7 | 4.7 | 7.2 | - | 1.2 | 1.1 | 8.3 |
| C17 :<br>1ω6c | - | - | - | - | - | - | - |
| C18 : 0 | 1.9 | 1.9 | 1.9 | 7.0 | 1.2 | 2.8 | - |
| C18 :<br>1ω7c | 15.7 | 8.5 | 12.2 | 6.1 | 3.7 | 14.9 | 10.5. |

The phylogenetic relationship of strain NG3^T^ was ascertained based on the 16S rRNA gene sequence similarity of NG3^T^ with other reported species using BLAST sequence similarity search (NCBI-BLAST/EzTaxon). The results revealed that at the 16S rRNA gene sequence level, strain NG3^T^ was close to the phylogenetic neighbours, *Rheinheimera aquatica* GR5^T^ (GQ168584) and *Rheinheimera texasensis* A62-14B^T^ (AY701891) with pairwise sequence similarity of 99% each. Phylogenetic analyses based on ML and NJ (Fig. 1) trees further indicated that strain NG3^T^ clustered with *Rheinheimera aquatica* GR5^T^ (GQ168584) and *Rheinheimera texasensis* A62-14B^T^(AY701891) with a phylogenetic distance of 99% each and distinct from the other species of the genus Rheinheimera. Despite the high 16S rRNA gene sequence similarity, it was observed that, at the whole genome level, when strain NG3^T^ was radioactively labelled, the DNA–DNA relatedness between strain NG3^T^ and *Rheinheimera aquatica* GR5^T^ (GQ168584), *Rheinheimera coerulea* TAPG2 (LT627667) and *Rheinheimera texasensis* A62-14B^T^(AY701891) is only 36, 34 and 32% (reciprocal reaction %) indicating that strain *Rheinheimera palamuruense* NG3^T^ is a novel species.

In addition, strain NG3^T^ could be phenotypically differentiated from the phylogenetically closely related species of *Rheinheimera aquatica* GR5^T^ (GQ168584), *Rheinheimera texasensis* A62-14B^T^(AY701891), *Rheinheimera chironomi* K19414^T^ (DQ298025), *Rheinheimera soli* BD-d46^T^ (EF575565) and *Rheinheimera tangshanensis* JA3-B52^T^ (DQ874340) (Tables 2). For instance strain NG3^T^ differs from both with respect to cell size, salt tolerance, growth temperature range, hydrolysis of starch and urea, citrate utilization, major fatty acid composition, cell wall and DNA G+C content (Table 2). It is interesting to see that the cell wall (peptidoglycan type) in two of the reference strains examined in the present study contained L-Orn–D-Glu which40 °C with optimum growth at 37°C. Growth was observed at salinities from 0 to 2% (NaCl, w/v), with an optimum between 0.7–1% (NaCl, w/v). Growth occurs from pH 6.5 to pH 8, with optimum growth at pH 7.0-7.5.

**Table 2.** Differential characteristics *of R. palamurensis* and related Rheinheimera species strains: 1, *Rheinheimera aquatica* GR5^T^ (GQ168584); 2, *Rheinheimera texasensis* A62-14BT(AY701891); 3, *Rheinheimera chironomi* K19414T (DQ298025); 4 *Rheinheimera soli* BD-d46T (EF575565); 5, *Rheinheimera tangshanensis* JA3-B52T (DQ874340).

| Characteristics | *Rheinheimera* sp. NG3T (GU566360) | *Rheinheimera aquatica* GR5<sup>T</sup> (GQ168584) | *Rheinheimera racoerulea* TAPG2 | *Rheinheimera aquatica* GR5<sup>T</sup> (GQ168584) | *Rheinheimera texasensis* A62-14BT (AY701891) | *Rheinheimera chironomi* K19414T (DQ298025) | *Rheinheimera soli* BD-d46T (EF575565) | *Rheinheimera tangshanensis* JA3-B52T (DQ874340) |
| --- | --- | --- | --- | --- | --- | --- | --- | --- |
| Source of isolation | Drinking water | Fresh water pond | Creek water | Fresh water pond | Lake water | Chironomid egg | Soil | Rice roots |
| Pigmentation | None | Greenish yellow | Creem to dark green | Greenish yellow | None | None | None | None |
| Cell width (mm) | 0.7-0.9 | 0.5-1.0 | 0.6-0.8 | 0.5-1.0 | 0.7-0.8 | 0.3-0.7 | 0.9-1.5 | 0.4-1.0 |
| Cell length (mm) | 1.1-2.0 | 1.5-2.0 | 1.0-1.2µm | 1.5-2.0 | 1.5-2.5 | 1.0-2.4 | 2.0-3.5 | 1.3-2.5 |
| Flagella* | S, P | S, P | S,P | S, P | S, P | S, P | S, P | S, P/L |
| Oxygen requirement | Aerobic | Aerobic | Aerobic | Aerobic | FAN | Aerobic | FAN | Aerobic |
| Growth temperature(°C) | 18-37° C | 10-40° C | 15-37° C | 10-40° C | 25-37° C | 4-40° C | 15-35° C | 10-37° C |
| Optimum | 27-32 | 35 | 25-30 | 35 | 30-37 | 30-37 | 25-30 | 30 |
| Growth in NaCl(% , w/v) | 0-2 | 0-0.2 | 0-1 | 0-0.2 | 0-1 | 0-2 | 0-4 | 0-3 |
| Optimum | 0.7-1.0 | 0.5-1.0 | 0.5 | 0.5-1.0 | 0.0 | 0.5-1.0 | 0-1.0 | 1.0 |
| pH for growth Range | 6.5-8.0 | 7.0-8.0 | 6.5-8.0 | 7.0-8.0 | 6.5-9.6 | 6.5-10.0 | 6.5-8.0 | 6.0-8.5 |
| Optimum | 7.0-7.5 | 8.0 | 7.0 | 8.0 | 7.5-8.0 | 7.5-8.0 | 7.0-7.5 | 7.0 |
| Nitrate reduction | + | + | - | + | + | + | + | - |
| Hydrolysis of aesculin | - | + | + | + | + | - | + | + |
| Assimilation of: D-Glucose | - | + | + | + | + | + | + | + |
| L-Arabinose | + | - | + | - | + | - | + | + |
| Mannitol | + | - | + | - | - | - | - | + |
| Maltose | + | + | + | + | - | + | + | + |
| Malate | - | - |  | - | - | + | - | + |
| Citrate | + | - | + | - | - | - | - | - |
| N-Acetylglucosamine | + | + | + | + | + | + | + | + |
| Hydrolysis of Tween 20 | + | + | + | + | + | + | + | + |
| Tween60 | - | + | + | + | - | - | - | - |
| Enzyme activity catalase | + | + | + | + | + | - | + | + |
| Esterase | - | - | - | - | - | - | - | - |
| Amylase | - | - | - | - | - | - | - | - |
| Majorquinones | MK-7, Q8 | Q8 | MK-7, Q8 | Q8 |  |  | MK-7, Q8 |  |
| DNA G+C content (mol%) | 48.5% | 51.9% | 53.6% | 51.9% | 48.2% | 49.9% | 49.2% | 47.0% |

On the basis of phylogenetic inference, strain NG3^T^ occupies a distinct position within the genus *Rheinheimera* that is supported by a unique combination of chemotaxonomic and biochemical characteristics. Strain NG3^T^ represents a novel species of the genus *Rheinheimera*, for which we propose the name *Rheinheimera palamurensis*sp. nov.

### Alignment with Bacteria, Fungi, Virus and Archaea genomes

The pre-processed reads were first aligned to the human genome (hg19) using BWA-MEMaligner to remove human genome contamination from the sample. The uncontaminated sequences were then taken for further alignment with known Bacterial, Fungal, Viral andArchaeal genomes using BWA-MEM aligner. Around 6.10 % of the reads map to the human genome, and 65 % to the bacterial genome.

### Reference guided Assembly

The human unaligned reads were aligned to the nearest reference genomeof species, which was predicted from the blast result. The reference fastafiles Rheinheimera sp. F8 were taken from NCBI and consensus was generated using the samtools tools version. Further, the coverage and depth statistics were generated using the bed tools version and in-house perl script.

### Phylogenetic tree

The 16s Sequence was extracted from Denovo assembled fasta and the extracted sequence was ran against NCBI nt blast. Here NODE_14 from the fasta was predicted to be the 16s sequence.

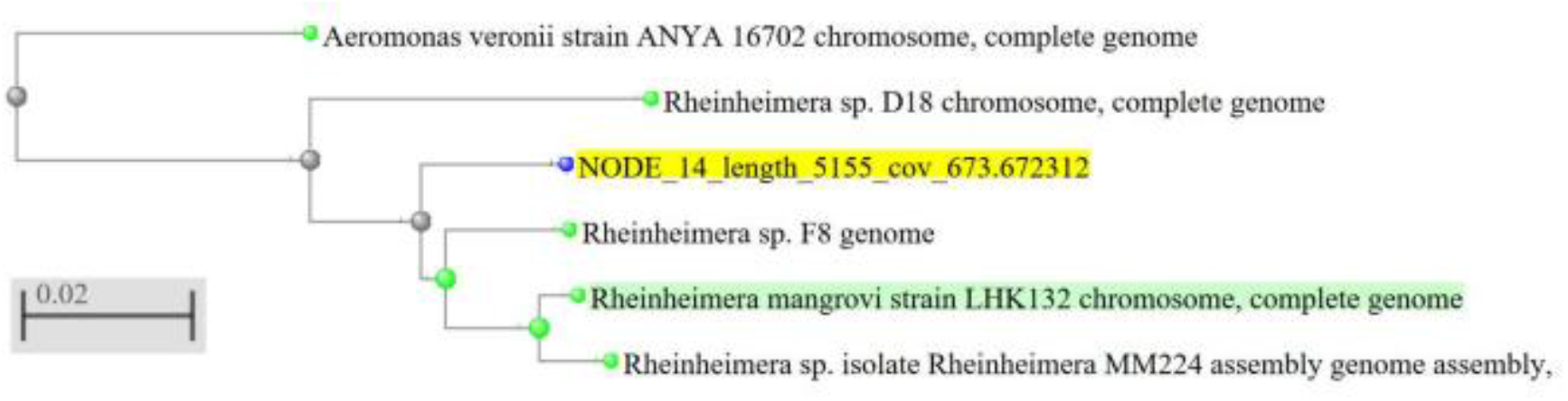

### Description of *Rheinheimera palamurensis* sp. nov

*Rheinheimera palamurensis* (pa.la.mu.ru.en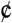se. N.L. neut. adj. palamurensis belonging to Palamuru). Cells are Gram-negative, aerobic, non pigmented and motile by means of a single polar flagellum. Growth occurred at 18 - 37° (optimum 27 - 32° C), pH at 6.5 – 8.0 (Optimum 7.0 – 7.5) and with 0-2% Nacl (Optimum 0.7-1.0%). Positive for Catalase, oxidase, phosphatase, lipase, nitrate reduction and urease but negative for esterase, amylase, protease and ornithine decarboxylase. Aesculin and Tween 80 are not hydrolysed. Assimilates Dextrose, Fructose, maltose, Galactose, Trehalose, Sucrose, L-arabinose, Mannose, Salicin, Glycerol and Inulin, ONPG but not Lactose, Xylose, Raffinose, Melibiose, Sodium gluconate, Dulcitol, Inositol, Sorbitol, Adonitol, Arabitol, Eythritol, α-Methyl-D-glucose, Rhamnose, Cellobiose, Melezitose, α-Methyl-D-mannoside, Xylitol, D-Arabinose, Malonate utilization and Sorbose. The isoprenoid quinone is MK-7 and Q8. The polar lipids comprise diphosphatidylglycerol, phosphatidylglycerol and phosphatidylethanolamine. The cell-wall peptidoglycan containsmeso-diaminopimelic acid as the diamino acid and cell-wall sugars are D-glucose and D-galactose. The genomic DNA G+C content of the type strain is 48.5 mol %, and the cellular fatty acid composition is given in Table 1.

The type strain is NG3^T^ (=GU566360=JCM-16716=KCTC-23111) was isolated from a drinking water sample collected from Nagar Kurnool, Telangana State, India.

## Acknowledgements

We thank Prof. G. N Srinivas, Vice-Chancellor, Palamuru University, Mahabubnagar, Telangana State, India for his encouragement and support.

## Funding information

Authors are grateful to DST (Department of Science and Technology, New Delhi, India) for the project **sanctioned vide no. *DST/TM /WTI/2k10/265 entitled “Advanced molecular methods for detection and identification of bacterial contaminants in potable water-purification and prevention”***.

## ‘Conflict of interest

All the authors read and approved the manuscript without any conflict before submission.

## Rheinheimera sp. NG3(2010) 16S ribosomal RNA gene, partial sequence

GenBank: GU566360.1

[GenBank Graphics](https://www.ncbi.nlm.nih.gov/nuccore/GU566360.1?report=graph)

~~~
>GU566360.1 Rheinheimera sp. NG3(2010) 16S ribosomal RNA gene, partial sequence
TTGCTCAGATTGAACGCTGGCGGCAGGCCTAACACATGCAAGTCGAGCGGGAACTTCGGTTCTAGCGGCG
GACGGGTGAGTAATGCGTAGGAAGCTACCCGATAGAGGGGGATACCAGTTGGAAACGACTGTTAATACCG
CATAATGTCTACGGACCAAAGTGTGGGACCTTCGGGCCACATGCTATCGGATGCGCCTACGTGGGATTAG
CTAGTTGGTGAGGTAATGGCTCACCAAGGCGACGATCCCTAGCTGGTTTGAGAGGATGATCAGCCACACT
GGAACTGAGACACGGTCCAGACTCCTACGGGAGGCAGCAGTGGGGAATATTGGACAATGGGCGCAAGCCT
GATCCAGCCATGCCGCGTGTGTGAAGAAGGCCTTCGGGTTGTAAAGCACTTTCAGTTGGGAGGAAGCGTT
GTGTGTTAATAGTACACAGCGTTGACGTTACCAACAGAAGAAGCACCGGCTAACTCTGTGCCAGCAGCCG
CGGTAATACAGAGGGTGCAAGCGTTAATCGGAATTACTGGGCGTAAAGCGCACGTAGGCGGTTTTTTAAG
TCAGATGTGAAAGCCCCGGGCTCAACCTGGGAATTGCATTTGAAACTGGAAAACTAGAGTGTGTGAGAGG
GGGGTAGAATTCCAAGTGTAGCGGTGAAATGCGTAGAGATTTGGAGGAATACCAGTGGCGAAGGCGGCCC
CCTGGCACAACACTGACGCTCAGGTGCGAAAGCGTGGGGAGCAAACAGGATTAGATACCCTGGTAGTCCA
CGCCGTAAACGATGTCTACTAGCTGTTCGTGACCTTGTGTCGTGAGTAGCGCAGCTAACGCACTAAGTAG
ACCGCCTGGGGAGTACGGTCGCAAGATTAAAACTCAAATGAATTGACGGGGGCCCGCACAAGCGGTGGAG
CATGTGGTTTAATTCGACGCAACGCGAAGAACCTTACCTACTCTTGACATCTACAGAAGACTGCAGAGAT
GCGGTTGTGCCTTCGGGAACTGTAAGACAGGTGCTGCATGGCTGTCGTCAGCTCGTGTTGTGAAATGTTG
GGTTAAGTCCCGCAACGAGCGCAACCCTTATCCTTAGTTGCCAGCACGTAATGGTGGGAACTCTAGGGAG
ACTGCCGGTGATAAACCGGAGGAAGGTGGGGACGACGTCAAGTCATCATGGCCCTTACGAGTAGGGCTAC
ACACGTGCTACAATGGTATGTACAGAGGGAGGCAAGCTGGCGACAGTGAGCGGATCTCTTAAAGCATATC
GTAGTCCGGATCGCAGTCTGCAACTCGACTGCGTGAAGTCGGAATCGCTAGTAATCGCAAATCAGAATGT
TGCGGTGAATACGTTCCCGGGCCTTGTACACACCGCCCGTCACACCATGGGAGTGGGTTGCAAAAGAAGT
AGGTAGCTTAACCTTCGGGAGGGCGCTTACCACTTTGTGATTCATGACTGGGGTGAAGTCGTAACAAGGT
AACCGTAGGGGAACCTGCGGCTGGA
~~~

